# Aerobic heterotrophy and RuBisCO-mediated CO_2_ metabolism in marine *Thaumarchaeota*

**DOI:** 10.1101/2019.12.22.886556

**Authors:** Linta Reji, Christopher A. Francis

## Abstract

*Thaumarchaeota* constitute an abundant and ubiquitous phylum of Archaea that play critical roles in the global nitrogen and carbon cycles. Most well-characterized members of the phylum are chemolithoautotrophic ammonia-oxidizing archaea (AOA), which comprise up to 5 and 20 % of the total single-celled life in soil and marine systems, respectively. Using two high-quality metagenome-assembled genomes (MAGs), here we describe a divergent marine thaumarchaeal clade that is devoid of the ammonia-oxidation machinery and the AOA-specific carbon-fixation pathway. Phylogenomic analyses placed these genomes within the uncultivated and largely understudied marine pSL12-like thaumarchaeal clade. The predominant mode of nutrient acquisition appears to be aerobic heterotrophy, evidenced by the presence of respiratory complexes and various organic carbon degradation pathways. Unexpectedly, both genomes encoded a form III RuBisCO. Genomic composition of the MAGs is consistent with the role of RuBisCO in nucleotide salvage, as has been proposed previously for archaea harboring the form III variant. Metabolic reconstructions revealed a complete nonoxidative pentose phosphate pathway (PPP) and gluconeogenesis, which can cyclize the RuBisCO-mediated carbon metabolic pathway. We conclude that these genomes represent a hitherto unrecognized evolutionary link between predominantly anaerobic basal thaumarchaeal lineages and mesophilic marine AOA, with important implications for diversification within the phylum *Thaumarchaeota*.

## Introduction

Archaea of the phylum *Thaumarchaeota* are among the most abundant microorganisms on the planet, constituting up to 20 % of single-celled life in marine systems alone (1). Although most characterized members of *Thaumarchaeota* are ammonia-oxidizing archaea (AOA), the phylum also encompasses several archaeal clades for which ammonia oxidation has not yet been demonstrated (e.g., Group 1.1c, and Group 1.3) (ref. 2). These basal, non-AOA members of the phylum have primarily been described in terrestrial systems such as anoxic peat soils (3), subsurface aquifer sediments (4), geothermal springs (5,6) and acidic forest soil (7). Availability of molecular oxygen on Earth is hypothesized to have influenced the evolution and habitat expansion of AOA from the basal anaerobic guilds (8).

A deeply-branching marine thaumarchaeal clade that has eluded cultivation and genomic analysis efforts is the pSL12-like group, also referred to as Group 1A or ALOHA group. First detected by DeLong et al., (2006; ref. 9) in the North Pacific Subtropical Gyre at station ALOHA, this clade appeared to be divergent from Marine Group 1 (MG1) AOA, clustering with a hot spring-associated crenarchaeal 16S rRNA sequence pSL12 (10). Mincer et al. (2007; ref. 11) suggested that at least some members of the clade may harbor the ammonia oxidation machinery, based on correlating abundances of the 16S rRNA gene and the *amoA* gene (*amoA* encodes the alpha-subunit of ammonia monooxygenase; conventionally used as the functional marker for AOA). The only available genomic information for the pSL12-like lineage comes from a fosmid clone library generated from the Mediterranean Sea (12). One of the three pSL12-like fosmid sequences recovered by Martin-Cuadrado and colleagues (2008; ref. 12) contained genes putatively involved in nitrogen fixation; however, there has not been genomic or biogeochemical evidence supporting this observation since. Several SSU rRNA gene surveys have detected the pSL12-like group in various marine systems such as the Atlantic Ocean (13), Mediterranean Sea (14), multiple Pacific Ocean transects (15), and the Northern Gulf of Mexico (16). Despite their suggested roles in N-cycle transformations, the metabolic adaptations of the pSL12-like lineage remain an open question.

Here we analyze the genomic repertoire and metabolic strategies of the pSL12-like lineage, based on two metagenome-assembled genomes (MAGs) obtained from seawater incubation metagenomes. In particular, we propose the existence of a form III RuBisCO-mediated CO_2_ fixation pathway in this clade, supporting heterotrophic growth on various carbon compounds. The high degree of phylogenetic and metabolic separation between these MAGs and typical marine thaumarchaeal clades suggests that the pSL12-like lineage represents an evolutionary link between anaerobic basal clades of *Thaumarchaeota* and aerobic marine ammonia-oxidizers.

## Materials and Methods

### Sample collection, incubation, and DNA extraction

Water column samples for AOA enrichment incubations were collected from Monterey Bay, CA, in May 2010. ASW2 was collected from 150 m at station M1 (36.747 N, −122.022 W), and ASW8 was collected from 200 m at station M2 further offshore (36.697 N, −122.378 W). After 8 years of incubation at 12 °C, 925 and 1000 mL each of the samples (for ASW2 and ASW8, respectively) were filtered using a 0.22 μm filter (Supor, Pall Inc, New York, USA). DNA was extracted using the DNeasy kit (Qiagen, Valencia, CA, USA), following the manufacturer’s protocol. To maximize DNA yield, DNeasy capture columns were eluted twice with 50 mL each of elution buffer, resulting in 100 mL total elution volume for each sample. DNA concentration was measured using Qubit Fluorometer (Invitrogen, NY, USA); 1.41 and 1.88 μg/ml DNA was obtained from ASW2 and ASW8, respectively.

### Metagenome sequencing, assembly and binning

Metagenome sequencing was performed as a part of a Community Science Program project with the DOE Joint Genome Institute (JGI); the samples were sequences (2 × 151 bp) using the HiSeq 2000 1TB platform. Read quality-filtering was carried out using the custom JGI script jgi_mga_meta_rqc.py (v2.0.0). Briefly, trimmed paired‐end reads filtered using BBDuk (17) (v37.50; BBTools software package, http://bbtools.jgi.doe.gov) were read‐corrected using BFC (v.r181; ref. 18). Reads without a mate pair were removed.

Quality-filtered reads were assembled using MEGAHIT (v1.1.3; ref. 19,20), using a range of k-mers (k=21,33,55,77,99,127). Contigs longer than 2000 bp were binned using two algorithms: MetaBAT2 (v2.12.1; ref. 21) and MaxBin2 (v2.2.6; ref. 22,23). Resulting bins were refined using the bin refinement module in metaWRAP (v1.2.2; ref. 24), and subsequently re-assembled using SPAdes (v3.13.0; ref. 25) to improve assembly quality. CheckM (v1.0.12; ref. 26) was used to assess bin completion. Taxonomic classifications were obtained using the GTDB-tk toolkit (v0.3.2; ref. 27). Dereplication based on average nucleotide identity was done using dRep (v2.3.2; ref. 28). Only bins with estimated completeness ≥70 % and contamination <10 % were retained for downstream analysis.

### MAG annotation and metabolic reconstruction

Prodigal (v2.6.3; ref. 29) was used for gene prediction, and functional annotations were obtained using Prokka (v1.13.7; ref. 30). Additionally, the BlastKOALA and GhostKOALA tool servers (31) were used to obtain KO annotations for genes predicted by Prodigal. KEGG-decoder (32) was used to estimate pathway completeness based on KO annotations, and the results were plotted in R (33). SEED annotations were obtained from the online Rapid Annotation using Subsystem Technology (RAST) server (34). Metabolic reconstructions were carried out using the ‘Reconstruct Pathway’ tool in KEGG mapper (https://www.genome.jp/kegg/mapper.html). TransportDB (v2.0; ref. 35) was used to predict membrane transporters; these annotations were further confirmed by BLASTp searches. SignalP-5.0 Server was used for signal peptide prediction (http://www.cbs.dtu.dk/services/SignalP-5.0/).

### Phylogenetic analyses

Anvi’o (v5.4; ref. 36) was used to compute a phylogenomic tree, based on a concatenated alignment of 30 ribosomal proteins obtained from the MAGs as well as selected reference genomes representing the known diversity within mesophilic *Thaumarchaeota*. MUSCLE (37) was used to generate the alignment. The final tree was computed using FastTree (38).

We used BLASTp (39) to search the MAGs for proteins of interest. Barrnap (v0.9; https://github.com/tseemann/barrnap) was used to identify ribosomal features. 16S rRNA sequences were aligned with reference sequences using MAFFT (40). RuBisCO reference sequences were obtained from Jaffe et al. (2009; ref. 41). Phylogenetic trees were computed using Mafft alignmnets in FastTree (38) with 1000 bootstrap replicates each. FastANI (42) was used to compute average nucleotide identity (ANI) between the MAGs.

### Assessing environmental distribution of MAGs

As part of the time-series microbiome survey in Monterey Bay, we previously obtained a depth-resolved dataset of 16S rRNA V4-V5 amplicon sequences, as well as metagenomes and metatranscriptomes (43,44). We were able to match one of the MAG-derived 16S rRNA sequences to an operational taxonomic unit (OTU) obtained in a time-series molecular survey targeting the V4-V5 region of the 16S rRNA genes. We estimated the relative abundance of this OTU as well as several others that clustered with sequences from the MAGs (these sequences had at least 90 % sequence identity).

We used three metagenome sets for read recruitment: (i) a depth- and time-resolved metagenome dataset from Monterey Bay; (ii) a North Atlantic Ocean depth profile from the TARA Oceans dataset; and (iii) a North Pacific Ocean depth profile from the TARA Oceans dataset. Bowtie2 (v2.3.5; ref. 45) was used to recruit metagenomic and metatranscriptomic reads against the MAGs. Read abundances were normalized as the number of reads mapping to kilobase of MAG per GB of metagenome (RPKG).

## Results and Discussion

The MAGs assembled here represent the first high-quality genomes reported for the pSL12-like lineage (completion estimates for the two MAGs are 88.8% and 97.08%, with < 3% contamination; Table 1). Their relative placement within the phylum *Thaumarchaeota* was confirmed by both phylogenomic and single-gene phylogenetic analyses (Fig. 1). Within a maximum-likelihood tree computed using a concatenated alignment of 30 conserved core ribosomal proteins, the two MAGs were placed as a sister-clade to all known ammonia-oxidizing *Thaumarchaeota* of Group 1.1a (marine AOA) and 1.1b (soil AOA) (Fig. 1a). Similarly, based on 16S rRNA gene phylogeny, the MAGs clustered with environmental clone sequences of the pSL12-like clade (Fig. 1b). The original hot spring pSL12 lineage (including the only available MAG for this lineage, DRTY7 bin_36, assembled from a hot spring metagenome; ref. 6) comprised a distant sister clade to the marine pSL12-like group.

**Table 1:** MAG statistics.

| MAG ID | Completion | Contamination | Number of<br>contigs | N50 | Number of<br>bases |
| --- | --- | --- | --- | --- | --- |
| ASW8_bin1 | 97.08 % | 2.912 % | 91 | 16957 | 996535 |
| ASW2_bin45 | 88.83 % | 0.97 % | 46 | 35482 | 918577 |

Estimates of genome completeness and contamination.

**Figure 1.**
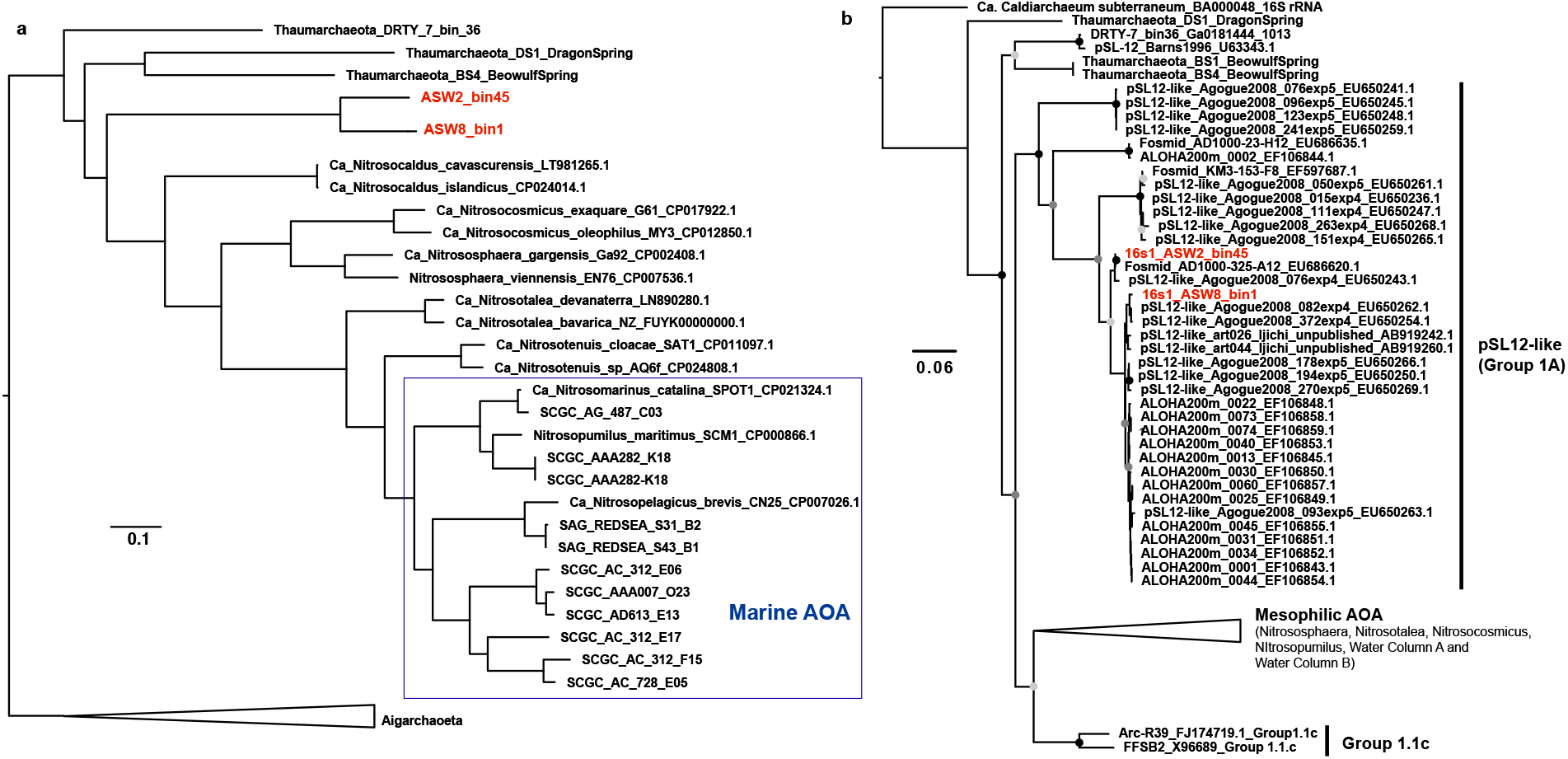
The assembled genomes cluster within the marine pSL12-like thaumarchaeal lineage. **a**, Phylogenomic tree computed using a concatenated alignment of 30 ribosomal proteins. **b**, Phylogeny of MAG-derived 16S rRNA gene sequences with genomic reference sequences.

### Metabolic potential distinct from typical marine Thaumarchaeota

Capacity for ammonia oxidation was not detected in either MAG, as we could not retrieve homologs of the ammonia monooxygenase (AMO) or nitrite reductase (*nirK*) genes. Moreover, the carbon-fixation pathway uniquely found in chemolithoautotrophic *Thaumarchaeota* - a modified version of the 3-hydroxypropionate/4-hydroxybutyrate (HP/HB) cycle (46) - appeared to be missing in both genomes. The myriad of copper-containing enzymes (e.g., multicopper oxidases, blue copper proteins) characteristic of AOA (47), were also missing. Since the genomes are not closed, our failure to detect these ‘expected’ pathways/genes does not definitively indicate their absence. However, there were striking differences in the overall genomic repertoire of typical AOA genomes and the MAGs recovered here (Fig. 2a), which cannot be explained by the lack of genome completeness alone.

**Figure 2:**
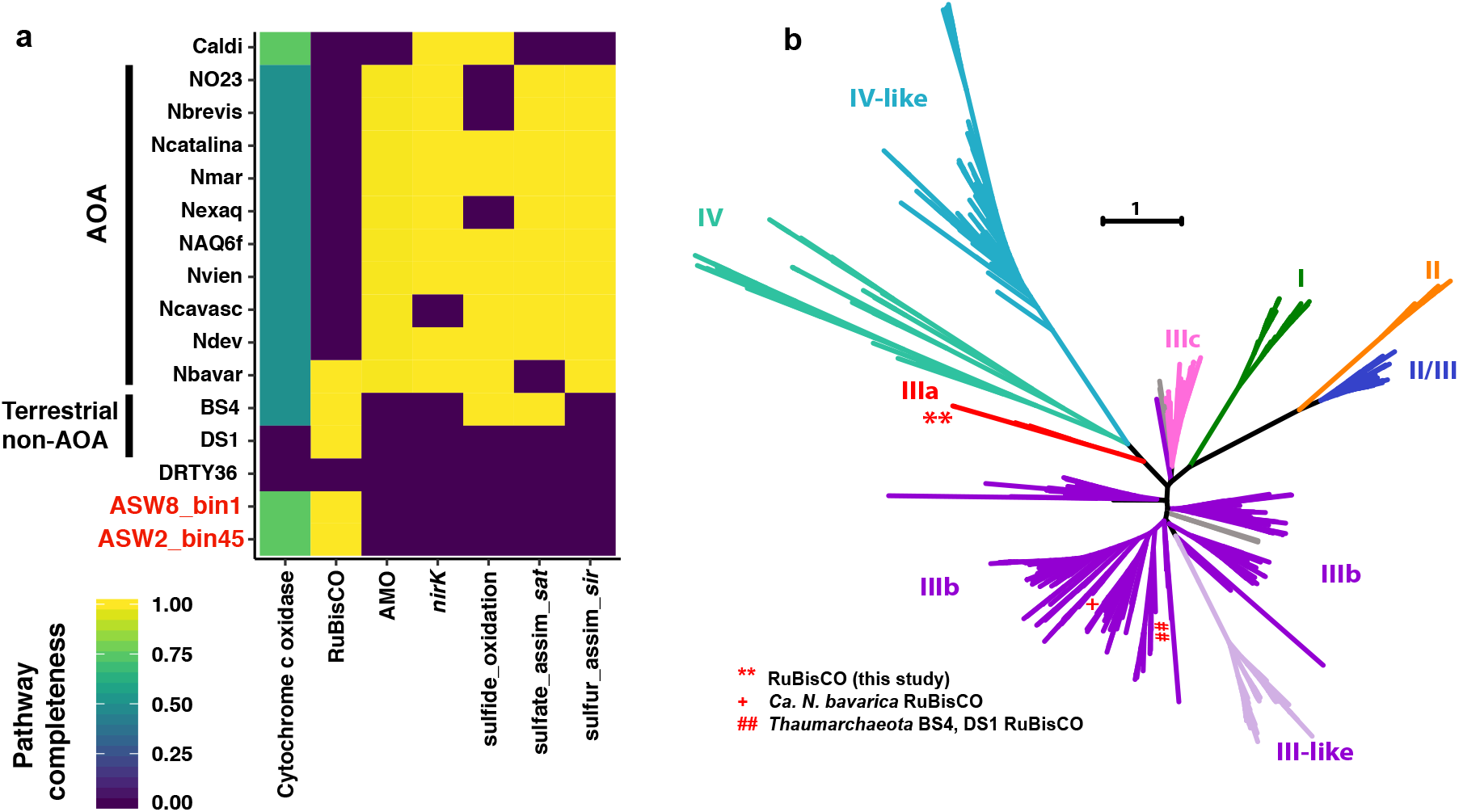
Metabolic capabilities of pSL12-like clade distinct from typical AOA. **a**, Comparison of selected metabolic pathways across thaumarchaeal genomes. pSL12-like MAGs are highlighted in red. *Caldiarchaeum subterraneum* belonging to the closely-related Phylum Aigarchaeota, is also included for comparison. Caldi: *Ca.* subterraneum; NO23: SCGC AAA007 O23; Nbrevis: *Ca.* Nitrosopelagicus brevis CN25; Ncatalina: *Ca.* Nitrosomarinus catalina SPOT01; Nexaq: *Ca.* Nitrosocosmicus exaquare; NAQ6f: *Ca.* Nitrosotenuis aquarius AQ6f; Nvien: *Nitrososphaera viennensis*; Ncavasc: *Ca.* Nitrosocaldus cavascurensis; Ndev: *Ca.* Nitrosotalea devanaterra; Nbavar: *Ca.* Nitrosotalea bavarica; BS4: *Thaumarchaeota* archaeon BS4 (MAG); DS1: *Thaumarchaeota* archaeon DS1 (MAG); and DRTY36: DRTY-7 bin_36 (MAG). **b**, Phylogenetic tree of RuBisCO sequences computed in FastTree using a MAFFT alignment of amino acid sequences. The MAG-derived RuBisCO sequences are highlighted. Previously reported thaumarchaeal RuBisCO sequences are also highlighted.

None of the six canonical carbon fixation pathways were complete in the MAGs. It is possible that these *Thaumarchaeota* may use the recently described reverse oxidative TCA cycle for CO_2_ fixation (48), since the genomes contained fumarate reductases, and 2-oxoglutarate/2-oxoacid ferredoxin oxidoreductases. In this pathway, a reversible citrate synthase catalyzes the production of citrate from acetyl CoA. Recently, metabolic reconstructions were used to predict the existence of the roTCA cycle in Aigarchaeota (6). We take caution in asserting roTCA CO_2_ fixation in pSL12-like *Thaumarchaeota*, since genomic inference alone is not sufficient evidence for this pathway (many of the enzymes are bifunctional and common with the anabolic TCA cycle).

The presence of respiratory complexes and various organic carbon-assimilating metabolic pathways (e.g., fatty acid oxidation, sugar metabolism, amino acid degradation and potential methylotrophy; Fig. 3) suggest that these *Thaumarchaeota* may be aerobic heterotrophs. The MAGs encoded several pyrroloquinoline quinone (PQQ)-dependent dehydrogenases containing N-terminal signal peptides (indicating extracellular localization), which can directly contribute reducing equivalents to the respiratory chain via extracellular sugar or alcohol oxidation (Fig. 3). Genome annotations suggest the potential for one-carbon (C1) compound utilization, particularly methanol and formaldehyde oxidation via a partial methylotrophic pathway. The PQQ-dependent methanol dehydrogenases likely oxidize methanol to formaldehyde and then to formate, using the tetrahydromethanopterin (H_4_MPT) route. The complete pathway could not be identified in either MAG; however, F420-dependent methylene-tetrahydromethanopterin dehydrogenases (*mtd*) were present in both genomes. The tetrahydrofolate (THF) pathway for formaldehyde and formate assimilation was complete in both MAGs.

**Figure 3:**
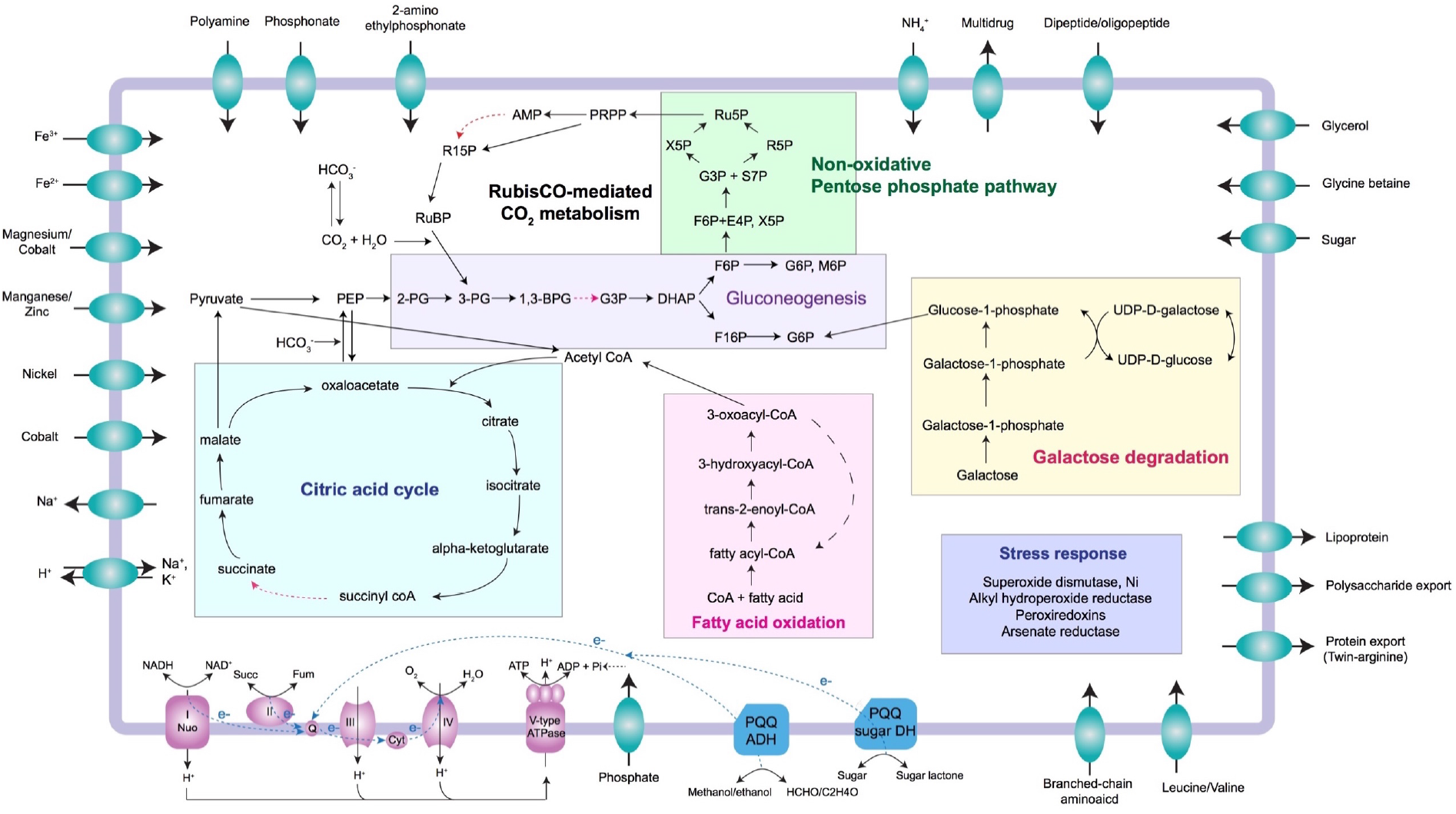
Overview of metabolic potential based on metabolic reconstructions of the pSL12-like MAGs. Red dashed arrows indicate unidentified genes. The TCA cycle is presented in the anabolic direction. For detailed gene information, see Supplementary Dataset 1.

Thaumarchaeal lineages previously identified as basal groups lacking the capacity to oxidize ammonia (which were obtained from non-marine environments) are reported to possess anaerobic energy generation pathways such as sulfate or nitrate reduction (5). The MAGs assembled here contained no evidence for anaerobic respiration. Moreover, many of the genomic features identified as unique/core features for the anaerobic basal thaumarchaeal lineages in a recent comparative meta-analysis (8) were also absent in these MAGs [(i.e., pyruvate:ferredoxin oxidoreductase (*porABDG*), cytochrome bd-type terminal oxidase (*cydA*), and acetyl-CoA decarbonylase/synthase (*codhAB*)]. Thus, multiple lines of evidence point to these MAGs representing a divergent, basal lineage within the aerobic, mesophilic clade of *Thaumarchaeota*.

### MAGs contain a methanogen-like form IIIa RuBisCO

Unexpectedly, both MAGs harbored an archaeal type III ribulose-bisphosphate carboxylase (RuBisCO) gene. Hypothesized to be the most ancient form of RuBisCO, form III is predominantly found in archaea (49). Recent surveys of metagenomic datasets have revealed numerous members of the candidate phyla radiation (CPR; ref. 50,51) and DPANN archaea (41,52) also encoding a form III-like RuBisCO. A divergent variant is found in methanogenic archaea, which is categorized as form III-a. Our MAG-derived sequences clustered with the methanogen III-a RuBisCO sequences (Fig. 2b), albeit with 30-35 % amino acid identity.

Two separate studies have previously reported a form III RuBisCO in *Thaumarchaeota*, and in both cases the assembled genomes represented acidophilic terrestrial lineages: (i) *Ca*. Nitrosotalea bavarica SbT1 was assembled and binned from an acidic peatland metagenome (53), and (ii) the deeply-branching strains BS4 and DS1 were assembled from acidic geothermal spring sediments in Yellowstone National Park (5). RuBisCO sequences from these MAGs clustered within the main archaeal form III clade (Fig. 2b), and were < 30 % identical (in the amino acid space) to the sequences we obtained in this study.

Despite exhibiting carboxylase activity in prior studies, genomic and biochemical evidence suggest that form III RuBisCO is not involved in carbon fixation via the canonical Calvin-Benson-Bassham (CBB) cycle (54,55). Regeneration of the RuBisCO substrate - ribulose 1,5-bisphophate (RuBP) - is a key step required for the cyclization of RuBisCO-mediated carbon fixation. In many archaea harboring RuBisCO, phosphoribulokinase (PRK) required for the regeneration of the RuBisCO substrate (RuBP) is missing (54), suggesting the absence of a functional CBB pathway. Intriguingly, methanogenic archaea harboring form III-a RuBisCO encode a PRK, yet are missing other key components of the CBB cycle (56). In light of these observations, two different pathways have been proposed for integrating RuBisCO-mediated CO_2_ fixation into the central carbon metabolism of form III-harboring microorganisms.

In one, methanogenic archaea possessing form III-a RuBisCO have been demonstrated to use the reductive-hexulose-phosphate (RHP) pathway for RuBP regeneration and, thus, RuBisCO-mediated carbon fixation (56). As demonstrated in *Methanospirillum hungatei*, RuBP regeneration in the RHP pathway involves the activity of PRK, which the organism encodes (56). In these methanogens, the ribulose monophosphate (RuMP) pathway (involved in methylotrophic formaldehyde assimilation and detoxification) is hypothesized to operate in reverse, fixing CO_2_ via RuBisCO and PRK. A key intermediate, fructose-6-phosphate, is derived from gluconeogenesis, which cyclizes the pathway (56).

While the RuBisCO sequences we retrieved from our MAGs resembled the form III-a methanogen RuBisCO (Fig. 2b), a PRK homolog could not be identified in either of the genomes. Furthermore, many of the key enzymes of the RHP and RuMP pathway were also absent, pointing to a different functional role for RuBisCO in these *Thaumarchaeota*.

The second proposed route for RuBisCO-mediated carbon metabolism involves nucleoside assimilation/ degradation via the archaeal AMP pathway (54,55). Briefly, adenosine monophosphate (AMP, retrieved from the phosphorylation of nucleosides) is converted to ribose 1,5-bisphosphate (R15P) by AMP phosphorylase (AMPase). Subsequently, R-15P is isomerized to ribulose 1,5-bisphosphate (RuBP) by ribose-1,5-bisphosphate isomerase (R15Pi). In an irreversible reaction, RuBisCO combines RuBP with CO_2_ and H_2_O to yield 3-phosphoglycerate (3-PG), which then enters the central carbon metabolism (via glycolysis or gluconeogenesis). Sato et al. (2007; ref. 54) proposed that the reductive pentose phosphate pathway, if present, may cyclize the above-described series of transformations, effectively rendering it a carbon-fixation pathway.

While two key enzymes of the AMP pathway - RuBisCO and R15Pi - could be identified in both MAGs, we could not detect an AMP phosphorylase (AMPase) homolog. However, even if the pSL12-like lineage lacks an AMPase, a modified version of the AMP pathway is still possible if R15P is generated from a compound other than AMP. The best candidate is phosphoribosyl pyrophosphate (PRPP), a key pentosphosphate intermediate in nucleotide biosynthesis. Both MAGs encoded a ribose-phosphate pyrophosphokinase, which forms PRPP from ribose 5-phosphate (R5P). PRPP is known to undergo abiotic disphosphorylation to yield R1,5-P (57). Alternatively, this reaction can be enzyme-mediated, most likely by a bifunctional Nudix hydrolase (58) (both MAGs contained a homolog for this gene). Thus, there potentially exists a direct route to R15P from PRPP, bypassing the requirement for an AMPase. The remaining transformations in the AMP pathway can follow as usual, generating 3-PG from RuBP. Intriguingly, both MAGs also encoded an adenine phosphoribosyltransferase, which converts PRPP to AMP. Given all of this, the archaeal AMP pathway (or a variant of it) is potentially operative in these *Thaumarchaeota*, which includes inputs from PRPP, and possibly AMP.

### Cyclization of the CO_2_-incorporation pathway via pentose phosphate pathway and gluconeogenesis

The complete set of genes participating in the non-oxidative branch of the pentose phosphate pathway (PPP) could be identified in both MAGs (i.e., ribulose 5-phosphate isomerase, ribulose 5-phosphate 3-epimerase, transaldolase and transketolase; Fig. 3). This pathway operating in reverse to generate R5P from gluconeogenesis intermediates, combined with the PRPP-(AMP)-RuBisCO transformations described above, might constitute a cyclic CO_2_ fixation pathway (54,59) in these *Thaumarchaeota* (Fig. 3). The overall pathway can therefore be summarized as: (i) R5P formation from fructose-6-phosphate via nonoxidative PPP; (ii) conversion of R5P to PRPP; (iii) abiotic or enzyme-mediated conversion of PRPP to R15P (potentially via AMP); (iv) formation of 3-PG via the RuBisCO-mediated carboxylation reaction; and (v) conversion of 3-PG back to fructose-6-phosphate via gluconeogenesis (Fig. 3, and Fig. S1). Several of the genes coding for key enzymes in the proposed pathway appeared to be colocalized on the same assembled contigs in both MAGs (Fig. S1), suggesting potential co-expression.

A gamma-class carbonic anhydrase (CA) was present in both genomes, which catalyzes the interconversion of CO_2_ and HCO_3_^−^. Unlike the CAs observed in terrestrial AOA, the pSL12-like CAs did not contain signal peptide sequences. This suggests its involvement in intracellular reversible dehydration of HCO_3_^−^ to CO_2_, facilitating CO_2_ incorporation via RuBisCO (or the roTCA cycle, if present).

### Distribution of the pSL12-like lineage in the water column

To assess the environmental distribution of the MAGs, we matched the MAG-derived 16S rRNA sequences to a previously generated 16S rRNA amplicon dataset from the Monterey Bay upwelling system (43). One of the MAG-derived 16S rRNA gene sequences (from ASW8_bin1) was an exact match to an operational taxonomic unit (OTU; #694), which comprised < 0.5% of the total thaumarchaeal abundance at any given time in the depths sampled. Three other OTUs were 90 % or more identical to the MAG-derived 16S rRNA sequences, but were much less abundant than OTU694. At any given time, this group of OTUs only comprised at most 0.5 % of thaumarchaeal abundance (Fig. 4a). As observed in previous surveys, this pSL12-like group of *Thaumarchaeota* appeared to be more abundant below the euphotic zone (11,13,15,16), with potential seasonal variations in relative abundances. Occasional abundance peaks were observed in the photic zone during spring at M1 (Fig. 4a), which likely reflects upwelled populations (station M1 is situated directly above the upwelling plume in Monterey Bay).

**Figure 4:**
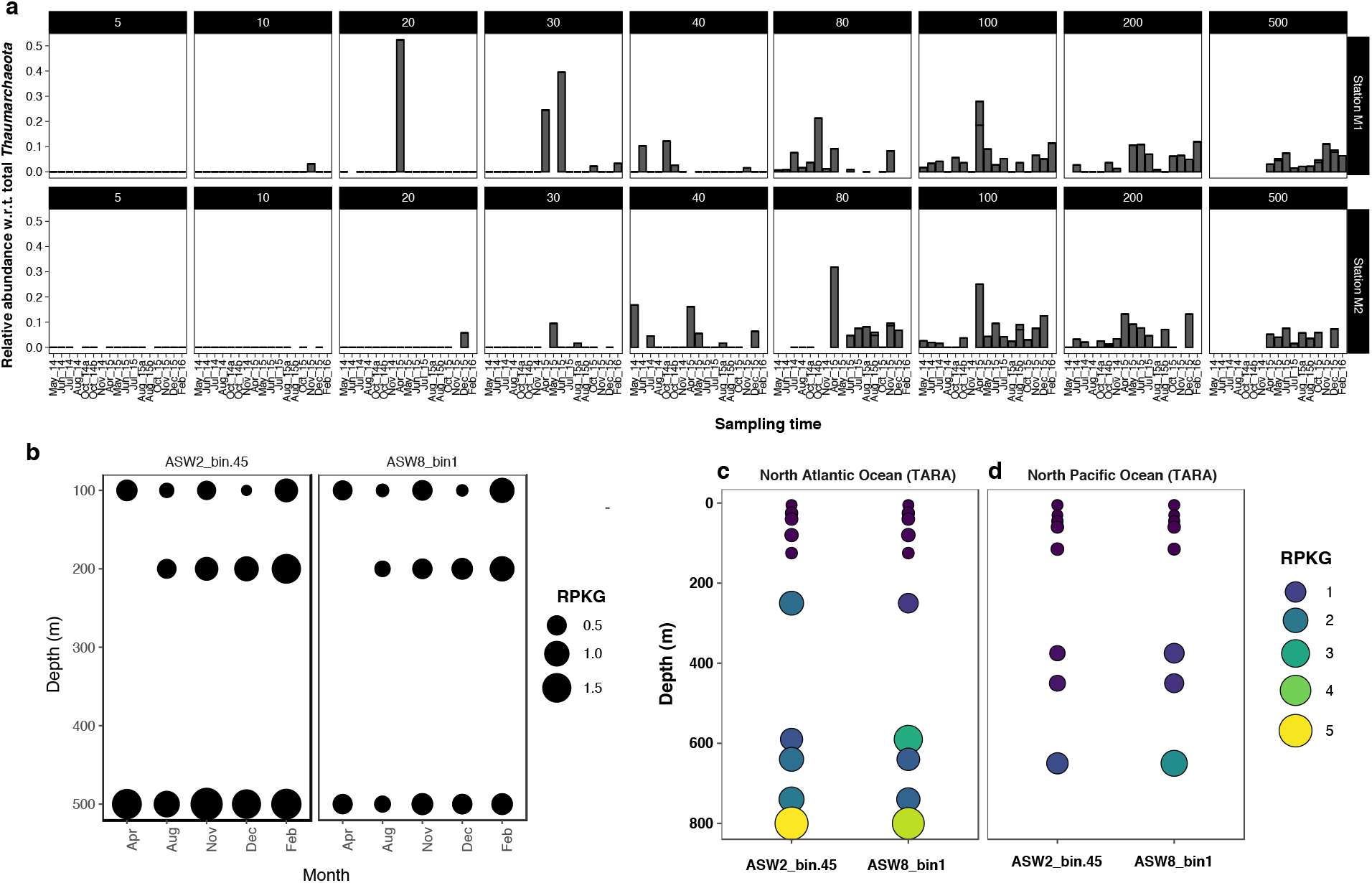
Distribution of pSL12-like lineage in Monterey Bay waters. **a**, Relative abundances (as a percentage of total thaumarchaeal abundance) of OTUs ≥ 90 % identical to the 16S rRNA gene sequences retrieved from the MAGs. The 2 major panels correspond to two sampling staions, M1 and M2, in Monterey Bay. Each subpanel represents a depth gradient between 5 - 500 m. **b**, Read recruitments of each MAG against Monterey Bay metagenomes. Size of the circle corresponds to normalized abundance. **c and d**, Metagenome read recruitments against Atlantic Ocean and Pacific Ocean depth profiles, respectively, from the TARA Oceans dataset. Relative abundances are presented as number of reads mapped per kilobases of genome per gigabases of metagenome (RPKG). Metagenome sample accessions are given in Table S1.

In recruiting metagenomic reads from Monterey Bay against the MAGs, we observed the highest recruitment at 500 m for ASW2_bin45. ASW8_bin1 recruited fewer reads, but appeared to have a relatively uniform abundance distribution across depths (Fig. 4b). Additionally, the relative abundances appear to change with seasonal hydrologic changes in the system (Fig. 4b). Recruitment against TARA Ocean metagenomes representing Atlantic Ocean and Pacific Ocean depth profiles revealed similar depth distribution of the pSL12-like lineage (Fig. 4b).

## Conclusions

In this work, we used reconstructed population genomes to infer metabolic adaptations of the elusive pSL12-like lineage of *Thaumarchaeota*, widely distributed in marine systems. The high-quality genomes described here offer a first glimpse into the genomic repertoire of a marine thaumarchaeal group devoid of an exclusively chemoautotrophic energy generation strategy. Only terrestrial basal lineages of *Thaumarchaeota* have been described thus far; the MAGs presented here represent the first genomic description of a basal lineage inhabiting the marine oxic environment. In this context, an especially intriguing consideration is the relative positioning of the pSL12-like clade within the thaumarchaeal evolutionary trajectory. These MAGs may help constrain the relative timing of the acquisition of aerobic metabolism and ammonia-oxidation within the phylum.

Overall, the divergent genomic features of the pSL12-like clade significantly alter our understanding of the metabolic diversity within this abundant archaeal phylum in the oceans. While further biochemical characterization is warranted to confirm the proposed metabolic transformations, our results suggest that obligate aerobic heterotrophy might be an overlooked metabolic strategy within pelagic *Thaumarchaeota*.

## Acknowledgments

Metagenome sequencing was carried out as part of a community science program (CSP) grant to C.A.F. from the DOE Joint Genome Institute. Computing for this project was performed on the Sherlock 2.0 cluster. We would like to thank Stanford University and the Stanford Research Computing Center for providing computational resources and support that contributed to the results presented here. We thank Marie Lund and Bradley B. Tolar for help with sample and data acquisition, respectively. We also thank Dr. Alfred Spormann for helpful feedback and discussion on an early draft of the manuscript. This study was supported (in part) by grant OCE-1357024 from NSF Biological Oceanography (to C.A.F.).

